# Deletion of VPS50 protein in mice brain impairs synaptic function and behavior

**DOI:** 10.1101/2023.07.04.547745

**Authors:** Constanza Ahumada-Marchant, Carlos Ancatén-Gonzalez, Henny Haensgen, Felipe Arancibia, Bastian Brauer, Rita Droste, H. Robert Horvitz, Martha Constantine-Paton, Gloria Arriagada, Andrés E Chávez, Fernando J Bustos

**Affiliations:** Instituto de Ciencias Biomedicas, Facultad de Medicina y Facultad de Ciencias de la Vida, Universidad Andres Bello, Santiago, Chile; Massachusetts Institute of Technology Cambridge, MA 02139, USA; Programa de Doctorado en Ciencias, Mención Neurociencia, Universidad de Valparaíso, Valparaíso, Chile; Instituto de Neurociencias, Centro Interdisciplinario de Neurociencia de Valparaíso (CINV), Facultad de Ciencias, Universidad de Valparaíso, Valparaíso, Chile

## Abstract

VPS50, is an accessory protein, involved in the synaptic and dense core vesicle acidification and its alterations produce behavioral changes in *C.elegans*. Here, we produce the mosaic knock out (mKO) of VPS50 using CRISPR/Cas9 system in both cortical cultured neurons and whole animals to evaluate the effect of VPS50 in regulating mammalian brain function and behavior. While mKO of VPS50 does not change the number of synaptic vesicles, it produces a mislocalization of the V-ATPase pump that likely impact in vesicle acidification and vesicle content to impair synaptic and neuronal activity in cultured neurons. In mice, mKO of VPS50 in the hippocampus, alter synaptic transmission and plasticity, and generated robust cognitive impairments associate to memory formation. We propose that VPS50 is an accessory protein that aids the correct recruitment of the V-ATPase pump to synaptic vesicles, thus having a crucial role controlling synaptic vesicle acidification and hence synaptic transmission.

## Introduction

Thousands of mutations have been implicated in developing neurodevelopmental disorders, each showing different degrees of certainty and producing a myriad of phenotypes ^1–6^. Thus, studying the specific gene mutations and their phenotype is crucial to finding common pathways and potential therapies for these disorders. Among neurodevelopmental disorders, autism spectrum disorders (ASD) are neurodevelopmental disorders characterized by deficits in social interaction, repetitive behaviors, and anxiety ^7^. These phenotypes are mainly produced by genetic mutations that cause changes in brain wiring, structure, and function. More than 1300 mutations have been associated with ASD, depicting the complexity of its multifactorial genetics and the phenotypes produced ^1–6^. Mutations that affect the function, filling, or availability of synaptic vesicles have been strongly associated with the appearance of ASD phenotypes ^8^. For instance, mice carrying mutations in Syn1, a synaptic vesicle component causing limited synaptic vesicle release, or Nhe9, a Na/H exchanger that causes hyper-acidification of vesicles, show phenotypes associated with ASD ^9–11^. Recently, the function of VPS50 in *C. elegans* controlling behavioral states and its expression in murine culture neurons was described^12^. VPS50 is an accessory protein widely expressed in the nervous system starting early in embryonic life ^12^. Recent studies have demonstrated that VPS50 is associated with some components of the Golgi-associated retrograde protein (GARP) complex, and more specifically with the endosome-associated recycling protein (EARP) complex, suggesting a role in endocytosis of synaptic vesicles by early endosomes ^13,14^. VPS50 physically interacts with the protein VHA-15, a component of the V-ATPase complex pump responsible for acidifying both dense core and synaptic vesicles, and its knockdown in cultured neurons shows a robust decrease in synaptic vesicle acidification ^12^. In humans, a deletion spanning the *Vps50* human gene, and the calcitonin receptor has been reported in an ASD patient ^15^. More recently, two individuals with homozygous loss of function mutations for *Vps50* have been described with a severe neurodevelopmental disorder ^16^. These findings underscore the potential role of VPS50 in ASD. However, no defined mechanisms have been identified that could explain the phenotypes shown by these individuals.

Here we used the CRISPR/Cas9 system to produce the mosaic KO (mKO) of VPS50 in cultured neurons and animals. We determined the effects on synaptic function, to later analyze mouse behavior to determine VPS50’s association to ASD, and in particular cognitive impairment.

Our findings provide insights into the role of VPS50 in synaptic function and behavior, as well as elucidate the mechanisms through which VPS50 mutants can contribute to ASD phenotypes. The brain mosaic VPR50 KO animal model will enable us to further investigate the cellular and molecular mechanisms underlying the function of VPS50 and its implications in ASD.

## Materials and Methods

### Animals and injections

All animal procedures and experiments were performed according to the NIH and ARRIVE guidelines and were approved by the animal ethics committee from Universidad Andrés Bello (020/2018). Newborn Cas9 KI mice (C57BL/6J; JAX 026179) were cryoanesthetized in a cold aluminum plate and injected with 1 μL of concentrated AAV (1×10^11^ vg) in each cerebral ventricle at a depth of 3 mm in the animal’s head at 2/5 of the intersection between lambda and the eye with a 10 μL HAMILTON syringe (Hamilton, 7653-01) and a 32 G needle (Hamilton, 7803-04). After de injection, P0 mice were placed in a heating pad until they recovered their color and temperature, then they were returned to their cage with the mother ^17–19^. Control mice were the same age as the injected ones. In the third week after birth, mice from both conditions were weaned off and separated by sex in cages with a 12/12 light/dark cycle with free access to food and water. A chip (p-chips, Pharmseq) was put in the tail of each animal for easy tracking during behavioral testing.

### Neuronal Cultures

At P1 neonatal mice were quickly decapitated and dissected in ice cold HBSS to obtain cerebral cortices as in ^18–21^. Cortices were minced and incubated for 20 min at 37C with papain (Worthington, USA) for enzymatic digestion. Then cells were transferred to a 15ml tube containing plating media (D-MEM supplemented with 10% Fetal bovine serum and 100 U/ml penicillin/streptomycin (Life technologies 15070-063). Cells were resuspended by mechanical agitation through fire-polished glass Pasteur pipettes of decreasing diameters. Cells were counted and plated on freshly prepared poly-L-lysine-coated plates. 2 hours later plating media was replaced with growth media (Neurobasal-A (Life technologies 1088802) supplemented with B27 (Life technologies 17504044), 2 mM L-glutamine (Life technologies 25030-081), 100 U/ml penicillin/streptomycin (Life technologies 15070-063)]. Half of the media was replaced every 3 days.

### AAV Production

AAV viral particles containing sgRNA directed to VPS50, red fluorescent protein tdTomato, hSyn promoter, and PHP.eB capsid ^22^ were obtained from HEK 293T cells and purified as described in^18,19^. To produce concentrated AAV viral particles with the plasmid containing the VPS50 sgRNA and the fluorescent protein tdTomato mediated by the hSyn promoter and with the PHP.eB capsid ^18,19,22^. In addition, AAVs were prepared using plasmids coding for ChR2 (Addgene Cat#28017), GCamP7f (Addgene Cat#104488), and custom plasmids coding for eCas9 ^23^, SyPhy ^24^, and pHoenix for synaptic vesicle acidification ^25^. HEK 293T cells were grown to approximately 6×10^4^ cells/cm^2^ with DMEM 10% FBS. Cultures were transfected using PEI “MAX” reagent (Polysciences, Cat 24765) with PHP.eB capsid plasmids, the vector with VPS50 sgRNA-tdTomato, and the helper plasmid DF6. After 24 h of transfection, the media was exchanged for DMEM 1% FBS. After 72h, media was collected from the plates and replaced with fresh DMEM 1% FBS. The collected media was stored at 4°C. 120 h after transfection, the cells were detached from the plate and transferred to 250 mL conical tubes, together with the collected media. They were centrifuged for 10 min at 2000 g, and the supernatant was removed and saved for later use. The pellet was resuspended in SAN digestion buffer (5 mL of 40 mM Tris, 500 mM NaCl and 2 mM MgCl2 pH 8.0) containing 100U/mL of Salt Active Nuclease (Arcticzymes, USA) and incubated at 37°C for 1 hour. The supernatant was precipitated using 8% PEG 8000 and 500mM NaCl. It was incubated on ice for 2 h and centrifuged at 4000 g for 30 min in 250 mL bottles. The supernatant was collected and resuspended with SAN digestion buffer. The solution was placed in an iodixanol gradient and ultracentrifuged at 350,000g for 2.5h. The phase containing the AAV was rescued and frozen at -80°C for later use.

### Brain sectioning and mounting

Half of the brains were submerged to assess brain infection and left to fix for a minimum of 24h in PBS CaMg + 4% PFA + 4% Sucrose into 30 mL flasks. After fixation, a Leica VT1000 vibratome was used to cut 100 μm coronal sections. Slices were kept in PBS CaMg 1X and mounted using Fluoromont G (EMS, Hatfield, PA) to preserve the fluorescence signal. Brain images were captured with a Nikon Eclipse TE2000 epifluorescence microscope.

### Protein extraction and electrophoresis

To carry out the immunodetection tests, proteins were extracted from the cortex and hippocampus of mice brains. 5 to 50mg of tissue was ground in N-PER lysis buffer (ThermoFisher Product No. 23225) with protease and phosphatase inhibitors (cOmplete^TM^ Protease Inhibitor Cocktail 11697498001, Roche Diagnostics) until a homogeneous solution was achieved and incubated for 10 min on ice. Then it was centrifuged at 10,000 g for 10 min at 4°C. Following the manufacturer’s recommendations, the supernatant with total proteins was collected and quantified using the BCA method (Perkin-Elmer). For electrophoresis, the proteins from the total extracts were denatured in loading buffer (NuPAGE LDS Sample buffer 4X; NP0007), heating at 95°C for 10 min. Then 40 μg of the sample was loaded on a 6% SDS acrylamide-bisacrylamide gel to visualize proteins larger than 100 kDa and 10% for proteins smaller than 100 kDa. Electrophoresis was carried out at 80-150 V constant voltage per gel in running buffer (25 mM Tris·Cl, 250 mM glycine, 0.1% SDS) in a minichamber (MiniPROTEAN System, BIO-RAD).

### Western Blot

After electrophoresis, the proteins were transferred to an activated PDVF membrane in transfer buffer (250 mM glycine, 25 mM Tris-Cl, 0.1% SDS, 20% methanol) at 400 mA constant current for 1.5 h. To verify protein transfer, the membrane was incubated with Ponceau red S (0.1% Ponceau S, 5% acetic acid). The membrane was washed with TBS/Tween 20 0.05% until the Ponceau red staining was removed. Next, the membrane was blocked for 1h to avoid non-specific protein binding sites in a 0.05% TBS/Tween 20 solution with 5% skim milk. The membrane was then incubated with the specific primary antibody VPS50 (Sigma, HPA026679-100UL, Rabbit, 1/500 dilution) in 0.05% TBS/Tween 20 solution with 5% skim milk at 4°C overnight. The membranes were washed with TBS/Tween 20 0.05% five times for 5 min each time. Then the membrane was incubated with a second antibody directed against the first antibody and coupled to horseradish peroxidase (HRP) in TBS/Tween 20 0.05% with 5% skim milk at room temperature for 1 h. Then, the membrane was washed with TBS/Tween 20 0.05% five times for 5 min each time. Detection was performed with chemiluminescence reagents (SignalFire^TM^ Elite ECL Reagent). Protein expression was normalized to β-Tubulin expression (Abcam Cat: ab6046, 1/500).

### DNA and RNA extraction

5 to 50mg of tissue was weighted for DNA and RNA extractions, and the Quick-DNA/RNA Miniprep kit (Cat: D7001) was used. Briefly, the tissue was homogenized with dounces in lysis buffer and centrifuged for 30s at 14,000g. The supernatant was collected and transferred to a Zymo-Spin™ IIICR column in a collection tube, centrifuged at 14,000g for 30s. The filtrate was used for RNA extraction and the column for DNA extraction. For RNA extraction, the same volume of 100% ethanol was added and homogenized, transferred to a Zymo-SpinTM IICR column, and centrifuged at 14,000g for 30s. The column was then treated with DNase I, and 400 μL DNA/RNA Prep Buffer was added, centrifuged, and washed twice with wash buffer. To elute the RNA, 25 μL of DNase/RNase-Free Water was added, incubated for 3 min, quantified, and stored at -80°C for later use. For DNA extraction, 400 μL DNA/RNA Prep Buffer was added to the Zymo-SpinTM IIICR column, centrifuged, and washed twice with wash buffer. To elute the DNA, 50 μL of DNase/RNase-Free Water was added, incubated for 5 min, quantified, and stored at -80°C for later use.

### RT-qPCR

RT-qPCR assays were carried out from 400 ng of extracted total RNA. To obtain complementary DNA (cDNA), each sample was mixed with 0.25 μg Oligo-dt (New England Biolab S1316S) in 10 μL to be denatured at 75°C for 5 min and then quickly transferred to 4°C for 5 min. Reverse transcription (RT) was performed in a final 20 μL volume containing: 10 μL denatured RNA, 100 U of M-MLV Reverse Transcriptase (NEB; M0253S), M-MLV RT buffer (NEB; B0253S) 1X, 20 U of RNase Inhibitor (NEB; M0314S) and 0.5 mM of dNTPs (Biotechnology N557-0.5ML). The mixture was incubated at 42°C for 1 hour, then at 95°C for 5 min, and the reaction was stopped at 4°C and then diluted five times with nuclease-free water. The cDNA was quantified by real-time PCR using 3 μL of the diluted RT mixture. Using relative abundance by the ddCt method, using the GAPDH gene as a loading control. Transcript detection was performed with specific primers for VPS50 mRNA (Sense: TGTTACTTCTCCGAGGCAGG, Antisense: GCTCTCAAAGGACCAAGAT) and GAPDH (Sense: ATGGTGAAGGTCGGTGTGAA, Antisense: CATTCTCGGCCTTGACTGTG).

### Calcium Imaging

Cortical neurons control and VPS50 mKO were co-infected at 3 DIV with GCaMP7f (Addgene Cat#104488). At 10 DIV, neurons were imaged using a Nikon Eclipse TE-2000 microscope equipped with a Co2/temperature chamber (Tokai-Hit). Pictures were acquired every 30ms for 5 minutes. The frequency of spiking was calculated using fluorescence over time plots using ImageJ.

### Electrophysiology cultured neurons

Whole-cell patch-clamp recordings on infected cortical neurons were performed and analyzed as previously described ^18,21,26^. The external solution contained (in mM) 150 NaCl, 5.4 KCl, 2.0 CaCl2, 2.0 MgCl2, 10 HEPES (pH 7.4), and 10 glucose. Patch electrodes (5–7 MΩ) were filled with (in mM) 120 CsCl, 10 BAPTA, 10 HEPES (pH 7.4), 4 MgCl2, and 2 ATP-Na2. After the formation of a high resistance seal and break-in (>1 GΩ), whole cell voltage and current signals were recorded with an Axopatch 700B amplifier (Molecular Devices). Signals were low pass filtered (5 kHz) and digitized (5–40 kHz) on a PC using pClamp 10 software. Cells were held at −60 mV.

To analyze synaptic function, we recorded isolated AMPA-mediated synaptic currents using a mixture of antagonists against NMDARs (20 μM d-APV) and GABAARs (2 μM bicuculline). In some experiments, the sodium channel blocker TTX (500 nM) was added to the bath to record miniature AMPA-mediated currents. For current-clamp analyses, cells were patched, and spontaneous spiking of cells was recorded. Depolarization of neurons was performed using square pulses or TTL activation of 488nm light to activate Channelrhodopsin channels. MiniAnalysis software (Synaptosoft) was used to analyze synaptic events during the tests. The frequency and amplitude of currents were automatically calculated and plotted.

### Hippocampal slice electrophysiology

Electrophysiological recording from hippocampal slices were conducted as previously described ^27,28^. Briefly, acute coronal hippocampal slices (400 mm thick) were prepared from control and VPS50 mKO at postnatal day (P) 30 to P45. Brain slices were cut using a DKT vibratome in a solution containing the following (in mM): 215 sucrose, 2.5 KCl, 26 NaHCO3, 1.6 NaH2PO4, 1 CaCl2, 4 MgCl2, 4 MgSO4, and 20 glucose. After thirty minutes recovery, slices were incubated in an artificial CSF (ACSF) recording solution containing the following (in mM): 124 NaCl, 2.5 KCl, 26 NaHCO3, 1 NaH2PO4, 2.5 CaCl2, 1.3 MgSO4, and 10 glucose equilibrated with 95% O2/5% CO2, pH 7.4. Slices were incubated in this solution for 30min before recordings.

All experiments, except where indicated, were performed at 28 ± 1°C in a submersion-type recording chamber perfused at 1–2 ml/min rate with ACSF supplemented with the GABA_A_ receptor antagonist picrotoxin (PTX; 100 mM). Extracellular field potentials (fEPSPs) were recorded with a patch pipette filled with 1mM NaCl and placed in the CA1 stratum radiatum. Whole-cell voltage-clamp recordings (Multiclamp 700B Molecular Devices, USA) were made from CA1 pyramidal neurons voltage-clamped at -60 mV using patch pipette electrodes (3–4 M) containing the following intracellular solution (in mM): 131 Cs-gluconate, 8 NaCl, 1 CaCl2, 10 EGTA, 10 glucose, 10 HEPES, pH 7.2, 292 mmol/kg osmolality.

fEPSPs and EPSCs were evoked by stimulating Schaffer collateral inputs with a monopolar electrode filled with ACSF and positioned ∼100–150 mm away from the recording pipette. Miniature EPSCs (mEPSCs) were recorded at 32 ± 1°C in the continuous present of tetrodotoxin (TTX, 500 nM) to block action potential dependent release, whereas spontaneous EPSC (sEPSCs) were recorded in the absent of TTX. Short-term synaptic plasticity was induced by two pulses (100 ms interstimulus interval) to calculate paired-pulse ratio (PPR) that was defined as the ratio of the slope or amplitude of the second EPSP/EPSC to the slope or amplitude of the first EPSP/EPSC, respectively. Long-term potentiation (LTP) was induced by 4 trains of 100 pulses at 100 Hz repeated four times, separated by 10 seconds. Reagents were obtained from Sigma, Tocris and Ascent Scientific, prepared in stock solutions (water or DMSO) and added to the ACSF as needed. Total DMSO in the ACSF was maintained less than 0.01%. For whole cells experiments, series resistance (range, 8–12 MW) was monitored throughout the experiment with a 5 mV, 80 ms voltage step, and cells that exhibited significant change in series resistance (20%) were excluded from analysis.

### Behavioral analyses

All behavioral tests on mice were carried out eight weeks after AAV injection and will be conducted as previously described ^18,19,21,29^. Before each test, mice cages were transported to the behavior room and habituated for 30 min in the dark. After completing a trial, the equipment and devices used were cleaned with 70% ethanol. Tests were recorded and analyzed with ANY-Maze software. Behavior tests were performed between 9:00 am and 6:00 pm. At the end of the battery of behavioral tests, the animals were sacrificed for subsequent molecular analyses.

#### Contextual fear conditioning

UGO-BASILE apparatus controlled by ANY-Maze was used. This equipment consists of a sound attenuating box, fan, light (visible/IR), a speaker, a USB camera, a single onboard controller, and a mouse cage. All trials were recorded, and all mice underwent habituation, conditioning, and testing phase. Twenty-four hours after training, the animals were tested for contextual memory. Each mouse was placed in the fear conditioning box, allowed to explore for 5 min freely, and returned to its cage. The number of freezing episodes and freezing time were registered.

#### Barnes Maze

A non-reflective gray circular platform (91 cm diameter) with 20 holes (5 cm diameter) evenly distributed along the perimeter, with one hole containing a metal escape tunnel, was used. Three exogenous visual cues (length/width ∼30 cm) were used around the platform: a black circle, a blue triangle, and a yellow square. The light was adjusted to 1000 lux in the center of the platform. All animals underwent a phase of habituation, spatial acquisition, and testing. On test day, the position of the escape tunnel was changed, and the animal was brought in the start box to the center of the platform, left for 10 s, and sound reproduction was started. The test ended at 90 seconds, or when the mouse found the escape tunnel. The number of primary and total errors, primary and total latency, and total distance before finding the gap were recorded. The number of visits to each hole was also measured to show preference.

### Data analysis

All values are presented as means ± standard error (SE) for three or more independent experiments. Statistical analyzes were performed using Student’s t-test. Values of p<0.05 are considered statistically significant. All statistical analyzes were performed using Graphpad Prism.

## Results

### VPS-50 regulates synaptic vesicle acidification and V-ATPase pump localization in cortical neurons

In *C. elegans*, VPS50 knock-out causes significant changes in behavior likely due to changes in synaptic and dense core vesicle acidification ^12^. In mice, VPS50 is highly expressed in the central nervous system and, its knock down in culture causes a deregulation of synaptic vesicle acidification ^12^. To further investigate the role of VPS50 in regulating mammalian synaptic function and behavior, we used CRISPR/Cas9 technology to get better insights into the mechanisms and consequences of VPS50 knock-out in mouse brain function. Cortical neurons from Cas9 KI animals were infected at 3 DIV with AAV coding for tdTomato as an infection marker alone or together with sgRNAs targeting *Vps50*. Transduction efficiency reached >90% of plated neuron in culture. Genomic DNA was extracted ten days after infection, and a surveyor assay was performed to determine the efficiency of gene edition. A combination of different sgRNAs targeting the *Vps50* genomic locus was tested, finding increased efficiency when pair of sgRNA1-6 was used (Supplementary Figure 1A). We performed the following experiments using sgRNA1-6 (hereafter VPS50 mKO) (Figure 1A). Edition of the locus causes a ∼70% reduction in VPS50 mRNA and protein levels in cortical neurons (Figure 1 B-D), confirming an efficient mKO of VPS50 after gene editing. Locus-specific sequencing shows the resulting edited genomic sequence caused by CRISPR/Cas9 (Supplementary Figure 1B). Importantly, VPS50 KO neurons shows no significant difference in the total number of synaptic vesicles assesses by electron microscopy (Figure 1E-F), however, we found a robust reduction in vesicle acidification assessed by ratio-SyPhy probe ^30^ (Figure 1G), consistent with the idea that VPS50 regulate synaptic vesicle acidification ^12^.

**Figure 1.**
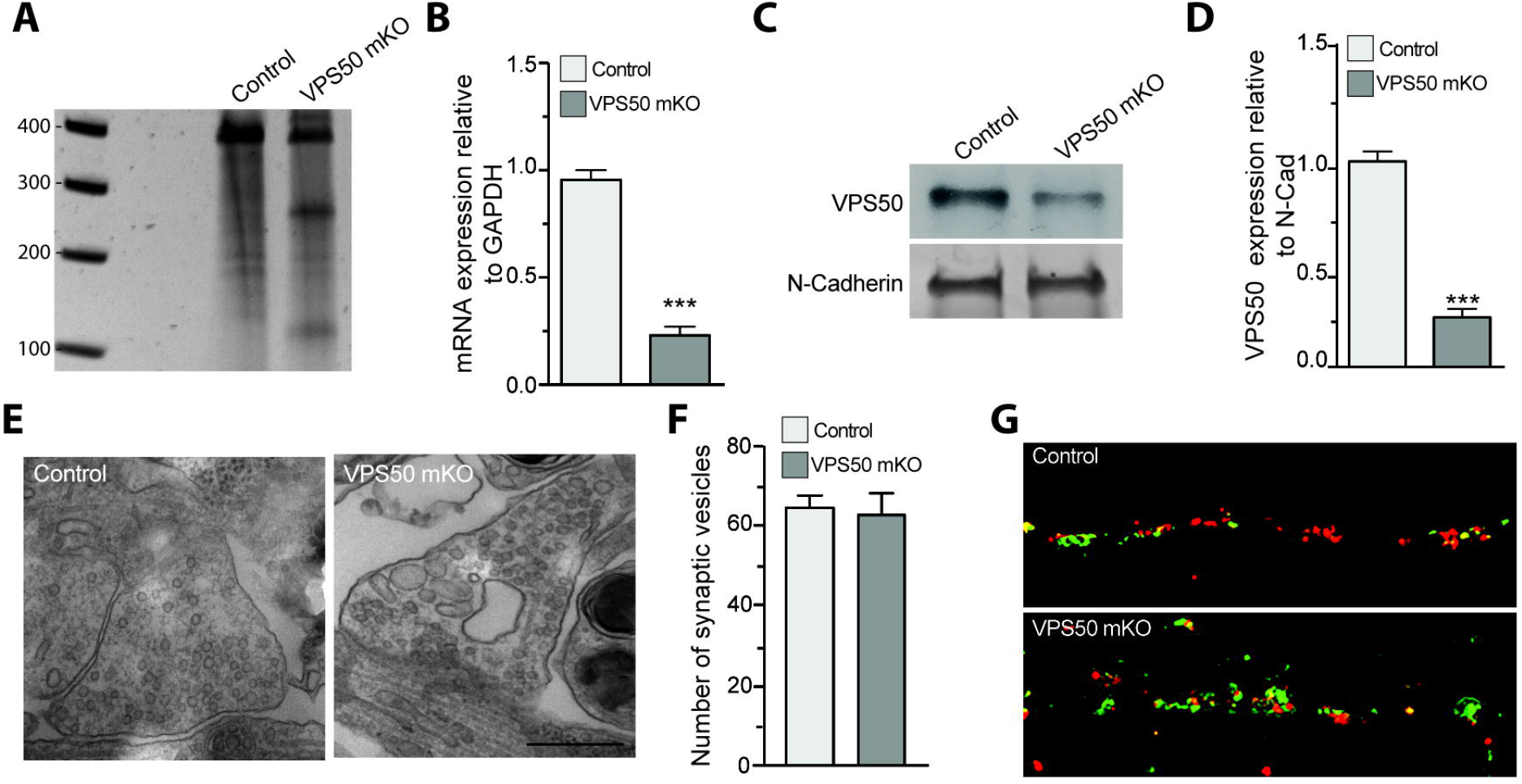
VPS50 gene edition causes decrease in synaptic vesicle acidification but no change in synaptic vesicle number. (A) Surveyor assay of Control or VPS50 mKO cortical neurons. (B) RT-qPCR to quantify relative mRNA expression of VPS50 mKO. (C-D) VPS50 protein expression in VPS50 mKO and Control neurons. (E-F) Representative images of synaptic terminals by electron microscopy and quantification of the number of synaptic vesicles in Control and VPS50 mKO neurons (n=50 cells per condition). (G) Representative images of Ratio-SyPhy signal to determine synaptic vesicle acidification in Control and VPS50 mKO neurons. Scale bar, 25um. ***p<0.001.

As VPS50 is enriched in synaptic and dense core vesicles as a soluble protein in mouse brain extracts ^12^, we next performed proximity ligation assays (PLA) to evaluate its approximate location within the synapse. First, we tested in control neurons whether VPS50 was close to the post-synaptic marker PSD95 or the presynaptic Synapsin1. We found that the PLA signal was only observed in VPS50/Synapsin1 condition, confirming that VPS50 is near synaptic vesicles (Figure 2A). As control, PLA reactions using Synapsin1/Synaptophysin and PSD95/Synapsin1 were used (Supplementary Figure 2). As VPS50 co-fractionates with the V-ATPase pump in mice tissue ^12^ and moreover, synaptic vesicle acidification is reduced in VPS50 mKO neurons (Figure 1D); it is possible that VPS50 might interact and/or help to localize the V-ATPase V1 pump to synaptic vesicles to acidify them for neurotransmitter filling. To test this possibility, first we use PLA to evaluate the proximity of VPS50 and V-ATPase pump in control neurons. We observed PLA signal indicating proximity between VPS50 and V-ATPase pump (Figure 2B). However, in VPS50 mKO neurons PLA signal is absent confirming the reduced expression of VPS50 and, consequently, the null interaction between VPS50 and V-ATPase pump (Figure 2B). Second, we evaluated whether the localization of V-ATPase pump was disrupted in VPS50 mKO neurons. While PLA experiments for the V-ATPase pump and Synapsin 1 (Syn) show that they are in proximity in control neurons (Figure 2C; top), no signal was observed in VPS50 mKO neurons (Figure 2C; bottom), strongly suggesting that knock-out of VPS50 causes a mis-localization of the V-ATPase pump in synaptic vesicle. Altogether, these results indicate that knockdown of VPS50 does not affect the total number of synaptic vesicles but produces a mis-localization of the V-ATPase pump that likely impair synaptic vesicle acidification and hereby synaptic function.

**Figure 2.**
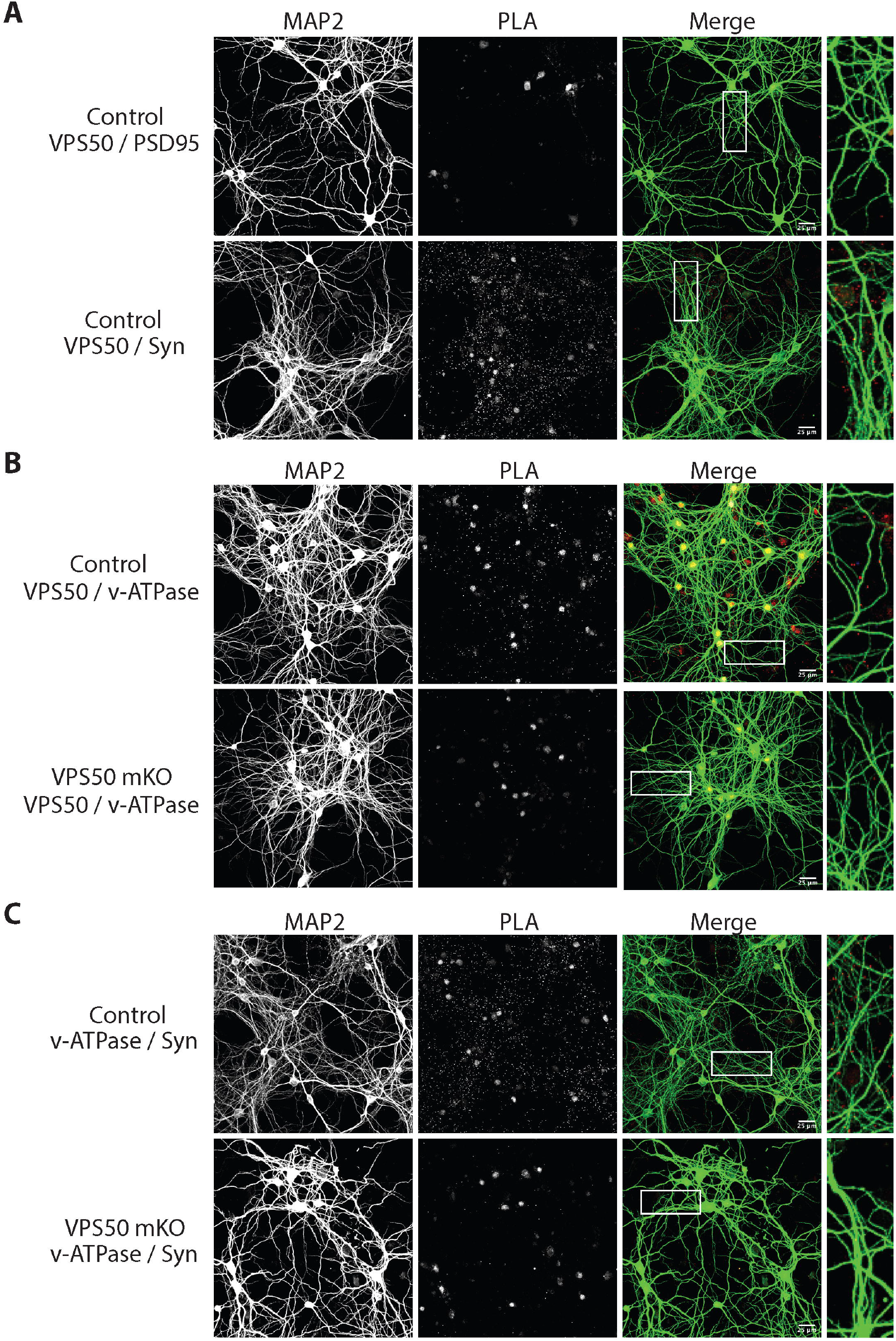
VPS50 mKO causes mislocalization of v-ATPase pump in cortical neurons. (A) PLA to determine pre- or post-synaptic localization of VPS50 in Control neurons using PSD95 or SynapsinI (Syn) antibodies. (B) Control and VPS50 mKO PLA to determine proximity of VPS50 and v-ATPAse pump. (C) PLA to determine localization of v-ATPase pump in Control and VPS50 mKO neurons. Inset shows magnification of the selected areas. Scale bar, 25 um.

### VPS50 mKO impairs synaptic activity in cortical neurons

To further evaluate the effect of VPS50 mKO in synaptic function, neurons were infected at 3 DIV and miniature and spontaneous excitatory postsynaptic currents (mEPSC and sEPSC, respectively) were recorded at 12-13 DIV. Compared to control neurons, VPS50 mKO neurons show a strong reduction in both the amplitude and frequency of sEPCS (Figure 3A-C). While the frequency and the amplitude of sEPSC does not change over 30-minute recordings in controlneurons, VPS50 mKO neurons display a significant reduction in the amplitude of the sEPSCs over time (Supplementary Figure 3A-C), consistent with the idea that vesicular content might be reduced in VSP50 mKO neurons. Moreover, VPS50 mKO neurons show a drastic reduction in the frequency but not in the amplitude of mEPSCs (Figure 3D-F), an effect that could reflect an increase in the exocytosis of empty or partially filled vesicles. Such reduction in the synaptic strength correlated with a decrease in the neuronal activity as the spontaneous spike frequency is decreased in VPS50 mKO neurons compared to control neurons (Figure 3G, H). In addition, calcium imaging using GCaMP7 shows that VPS50 mKO neurons have reduced calcium event frequency, confirming deficient spiking (Supplementary Figure 3D-E). To further evaluate if synaptic vesicle acidification is partly responsible for the reduction in vesicle content and neuronal activity, we used pHoenix, a genetically encoded proton pump targeted to synaptic vesicles ^25^. Spontaneous spiking of pHoenix infected control and VPS50 mKO neurons was recorded for 1 min as baseline, and then 532nm light was used to activate pHoenix. During the 2-minute activation period, we observed a partial recovery of spiking in VPS50 mKO neurons close to control levels (Figure 3 I-J; blue line). Moreover, after stimulation we observed no significant differences with control neurons (Figure 3K). Altogether, our results strongly suggest that the reduction in synaptic activity of VPS50 mKO requires synaptic vesicle acidification, a phenomenon that can be rescued by artificially acidifying synaptic vesicles.

**Figure 3.**
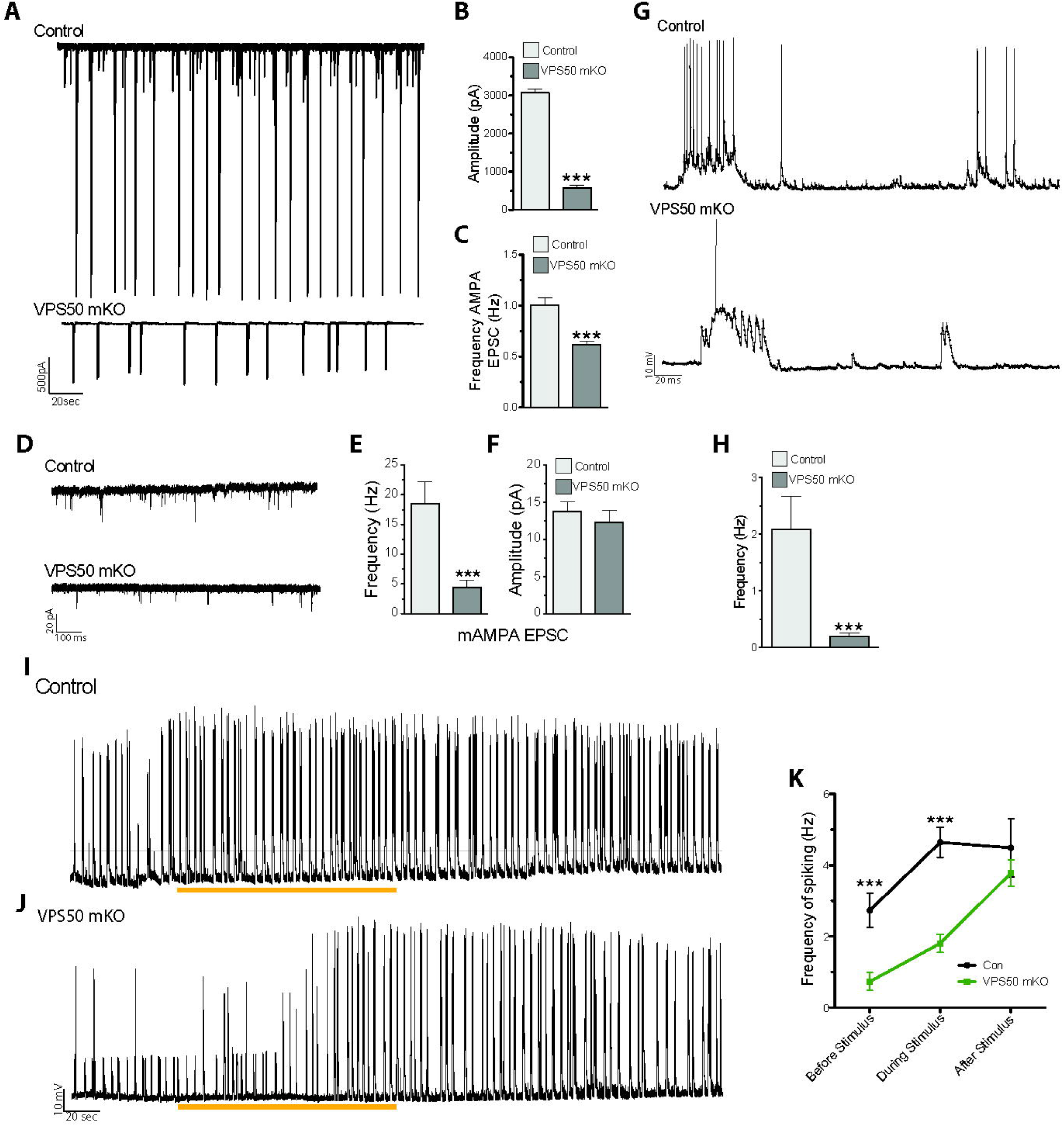
VPS50 mKO neurons show a reduction in synaptic activity that can be recovered by inducing synaptic vesicle acidification. (A) Representative traces and (B-C) quantification of Amplitude (B) and Frequency (C) of spontaneous AMPA-mediated EPSCs. (D) Representative traces and (E-F) quantification of the frequency (E) and amplitude (F) of miniature AMPA-mediated currents. (G) Current clamp of Control and VPS50 mKO neurons. (H) Quantification of the frequency of spiking in current clamp. (I-K) Current clamp representative traces of Control and VPS50 mKO neurons. Blue bars show the period pHoenix is activated. (K) Quantification of the frequency of spiking of Control and VPS50 mKO neurons before, during and after stimulus with pHoenix to acidify synaptic vesicles. At least 18 neurons from 3 independent experiments were analyzed. *** p<0.001.

### VPS50 mKO impacts synaptic function and memory formation

Once we have demonstrated that VPS50 mKO causes deficit in vesicle acidification and synaptic function in cultured cortical neurons, next we aimed to evaluate whether VPS50 mKO in mice might cause significant changes in behavior as suggested previously in *C.elegans* ^12^ and humans ^16^. Toward this end, we used the CRISPR/Cas9 system to induce VPS50 mKO in the mouse entire brain. We delivered high titer AAVs at postnatal day 1 (P1) intracerebroventricular in Cas9 KI animals, packed using the PHP.eB capsid ^22^ for its high efficiency in targeting the mouse brain. Animals were injected with the same sgRNAs tested in culture together with tdTomato or tdTomato alone as a fluorescence marker under the control of the human synapsin1 promoter. First, post-mortem analyses of animals show high AAV infection in multiples brain areas (Figure 4A), including the cortex and the hippocampus. In these brain areas, VPS50 protein expression is significantly reduced (Figure 4B), confirming an efficient mKO of VPS50 after gene editing in mouse brain. Second, we monitored basal synaptic function at Schaffer collateral to CA1 synapses in acute hippocampal slices. A significant decrease in the frequency but not in the amplitude of miniature excitatory postsynaptic currents (mEPSCs) was observed in VPS50 mKO synapses (Figure 5A-C). Likewise, a strong reduction in the frequency and amplitude of spontaneous EPSCs (sEPSCs) was also observed in the VPS50 mKO synapses (Figure 5D-F). Moreover, input/output curves revealed a large decrease in the amplitudes of evoked EPSC at all stimulus intensities tested in VPS50 mKO synapses (Figure 6A-B). Importantly, paired-pulse facilitation remains unchanged (Figure 6C-D) suggesting that the decrease in evoked EPSC amplitude cannot be accounted for changes in release probability, but could be due to a decrease in vesicle content and/or vesicle refilling by change the level of acidification as observed in cultured neurons (Figure 1G). To further evaluate whether the state of vesicle content might be involved in this synaptic change, we evaluated the response of VPS50 mKO synapses to high-frequency stimulation a condition in which one might detect the release of partially filled vesicles. Under this conditions, VSP50 mKO synapses response was greatly reduced compared to control synapses (Figure 6E-F), confirming that vesicle content is reduced in VSP50 mKO synapses. Next, we investigated the impact of VSP50 mKO in long-term synaptic plasticity (LTP) in the hippocampus, the cellular mechanism underlying learning and memory. We found that the magnitude of LTP induced by high-frequency stimulation was reduce in mice with disrupted VSP50 compared to control (Figure 6G-H), suggesting cognitive deficit in the context of memory formation in VPS50 mKO mice.

**Figure 4.**
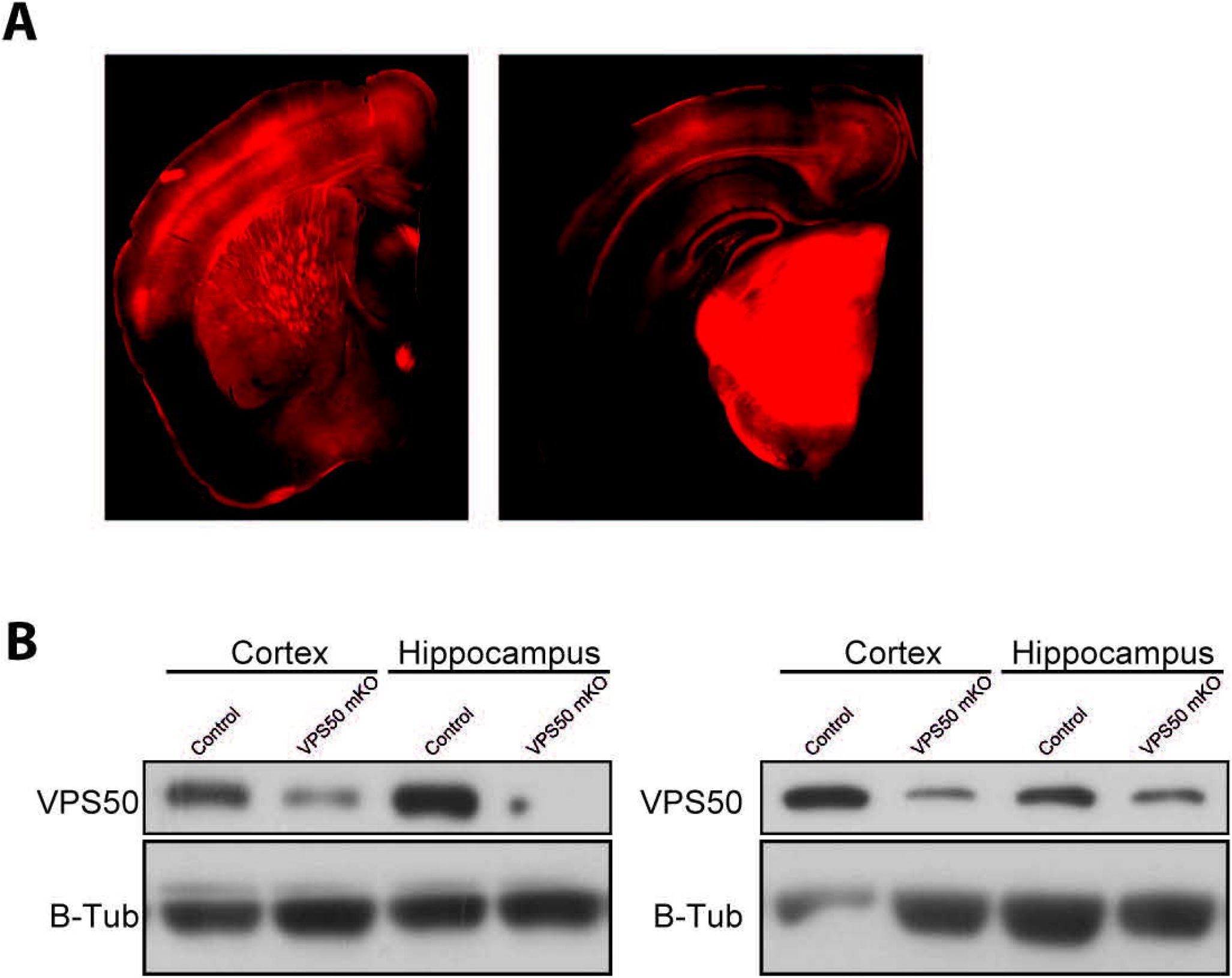
Systemic injection of AAV at P0 to produce VPS50 mKO. (A) Representative image of a coronal section of a mouse brain injected at P0 with AAV after 15 weeks. (B) Representative western blot analyses for the detection of VPS50 expression in Control and VPS50 mKO animals in both cortex and hippocampus. 2 animals are shown per condition. B-Tubulin is used as loading control.

**Figure 5.**
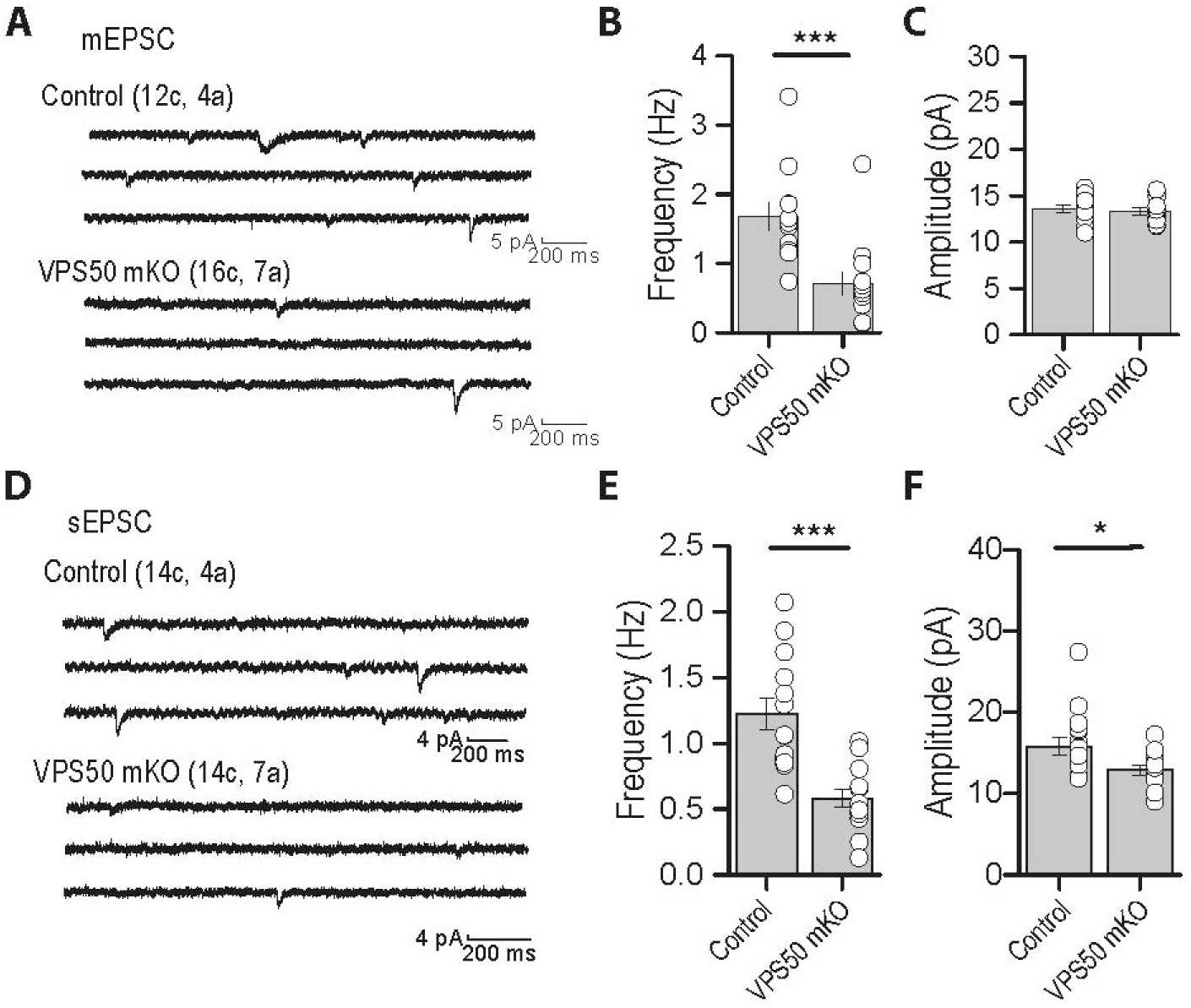
Hippocampal spontaneous excitatory synaptic activity is impaired in VSP50 mKO mouse. (A) Representative traces of mEPSC in hippocampal slices of Control and VPS50 mKO animals. (B-C) Quantification of the Frequency (B) and Amplitude (C) of mEPSC. (D) Representative traces of sEPSC in hippocampal slices of Control and VPS50 mKO animals. (E-F) Quantification of the Frequency (E) and Amplitude (F) of mEPSC. *p<0.05, *** p<0.001.

**Figure 6.**
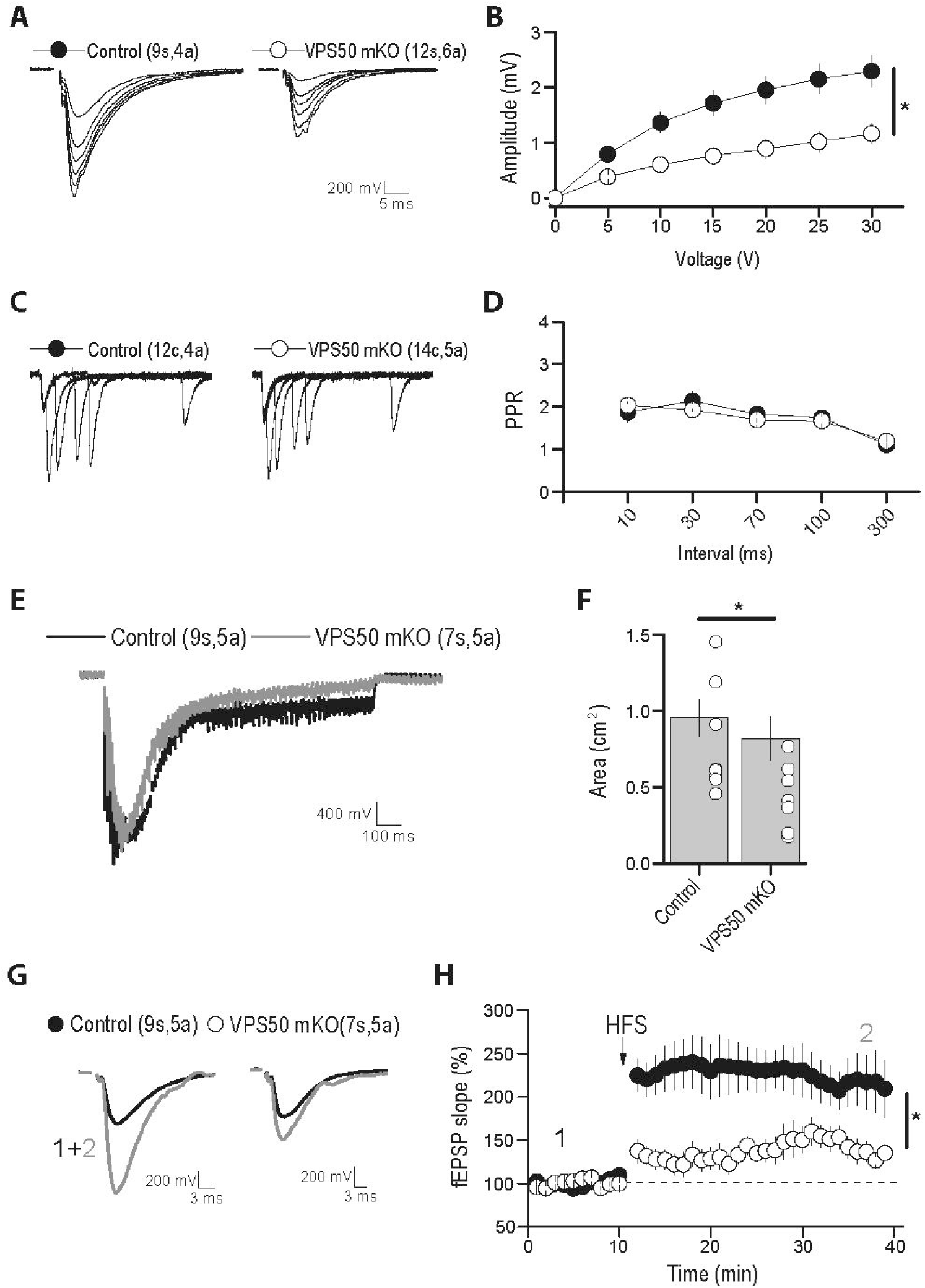
Hippocampal synaptic function and plasticity are impaired in VSP50 mKO mice. Electrophysiological recording of Control and VPS50 mKO hippocampal slices. (A) Representative traces of Input-output responses. (B) Input-output curves reveal a strong reduction in the amplitude of evoked EPSC at all stimulus intensity tested. (C) Representative traces and (D) quantification of paired-pulse facilitation at different inter-stimulus interval. (E) Representative traces and quantification (F) of the response to high-frequency stimulation. (G) Representative traces before and after LTP induction by high-frequency stimulation. (H) Quantification of potentials during LTP induction. Number of slices (s) or cells (c) and animals (a) are indicated in parenthesis. * p<0.05.

Finally, we evaluated VPS50 mKO mice performance in Barnes maze and Fear conditioning apparatus (Figure 6), two memory formation paradigms dependent on the hippocampus. First, we analyzed if VPS50 mKO mice model had locomotor problems that could affect their performance in behavioral testing. VPS50 mKO animals spent more time on the ramp in the accelerated rotarod apparatus than Control animals (Figure 7A), indicating that brain mKO of VPS50 have no impact on motor coordination. However, VPS50 mKO animals made a significantly higher number of primary errors (holes checked before finding the escape hole) on days 1 and 2 (Figure 7C), but behaved similar to control error numbers on days 3 and 4. We also found that primary latency, the time the animal takes to find the escape hole, was significantly affected in VPS50 mKO, but only on the first day (Figure 7D). Moreover, we found that VPS50 mKO animals spent significantly less time in the escape zone on days 1-3 compared to control littermates (Figure 7E). Lastly, using the context-dependent fear conditioning paradigm, we found that VSP50 mKO animals significantly decreased in freezing compared to control littermates (Figure 7F). Altogether, these results indicate that VPS50 mKO mouse display alteration in hippocampal synaptic function and memory deficits dependent on this brain area.

**Figure 7.**
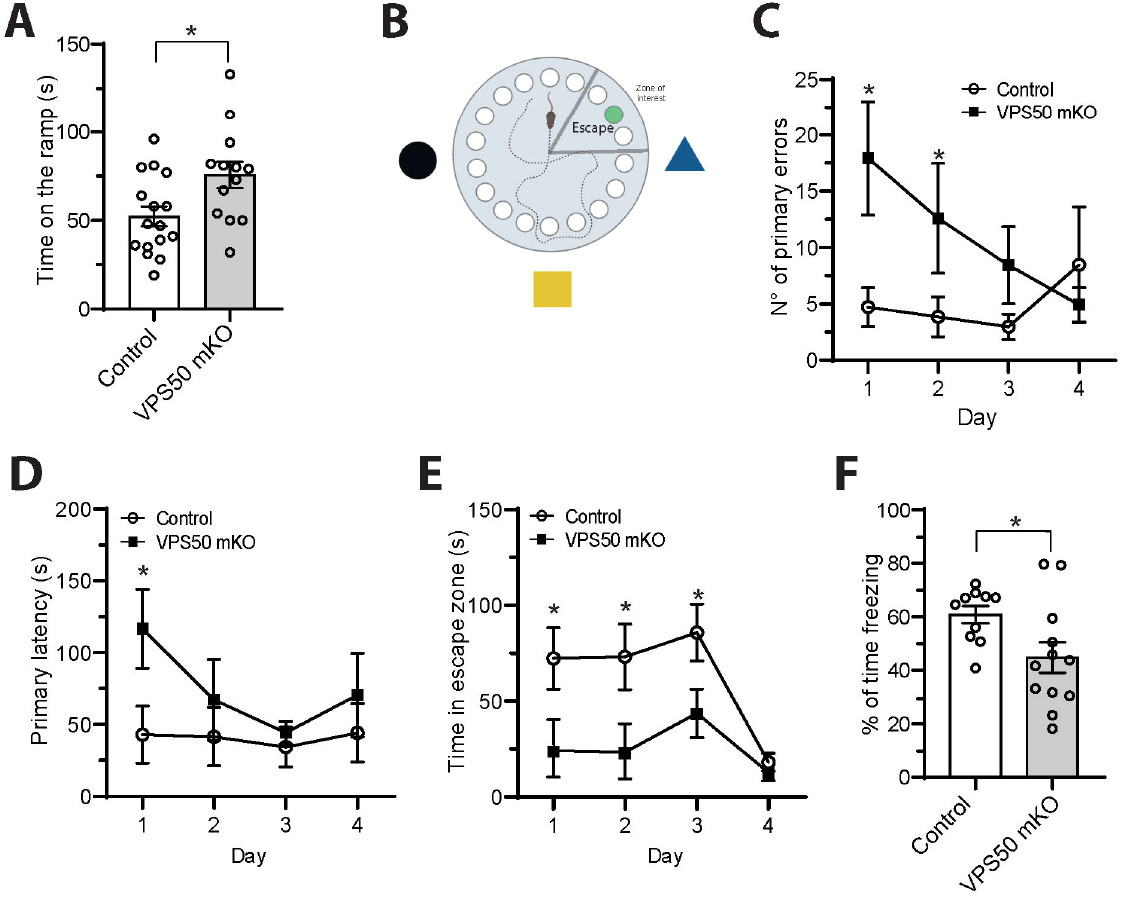
VPS50 mKO animals show impaired hippocampal memory formation. Brain wide Control and VPS50 mKO animals were subjected to behavioral testing. (A) Accelerated rotarod apparatus. (B) Scheme of the Barnes maze memory paradigm where clues attached to the wall, escape hole, and escape zone are shown. (C) Quantification of the numbers of primary errors before finding the escape hole. (D) Quantification of the primary latency to find the escape hole. (E) Quantification of the time spent in the escape zone. (F) Quantification of freezing percentage time in the fear conditioning paradigm. *p<0.05.

## Discussion

VPS50 is a conserved protein across multiple species including *C. elegans, M. musculus*, and humans, indicating its fundamental importance in cellular function ^12,16^. In the cell, VPS50 is associated to the EARP complex and more specifically to dense core and synaptic vesicles ^31^. Evidence from *C. elegans*, shows that VPS50 control locomotion behaviors and is associated to the V-ATPase pump ^12^.

The V-ATPase pump is recruited to synaptic vesicles after recycling serving as an initial step for neurotransmitters filling ^32^. Studies using *C. Elegans* show that worm mutants for unc-32, the homolog to the human V-ATPase pump, have severe neurotransmission deficits in motoneurons due to the lack of acidification and hence filling of neurotransmitters in their synaptic vesicles ^33^. Interestingly, the V-ATPase pump works by acidifying cellular compartments broadly in the cell, but some specificity is given due to the differential formation of the entire complex by different isoforms and proteins interactions ^34^.

Our findings reveal a critical role of VPS50 in regula synaptic function and behavior in mammals through its involvement in synaptic vesicle acidification. Using CRISPR/Cas9 to produce the mKO of VPS50, we discovered that VPS50 mKO neurons exhibit mislocalization of the V-ATPase pump, lacking proximity to synaptic vesicles, without affecting the total number of synaptic vesicles. Instead, this mislocalization affects vesicle acidification, thereby influencing vesicular content and/or vesicle refilling. Consequently, the limited acidification of synaptic vesicles leads to a significant reduction in neurotransmitter filling and a subsequent decline in synaptic activity ^8,32,35^. These findings strongly indicate that VPS50 plays a crucial role in facilitating the proper localization of the V-ATPase pump near synaptic vesicles to allow synaptic vesicle acidification. However, it remains to be investigated whether by the mislocated v-ATP pump the extent of vesicle filling is impacted.

Furthermore, our data demonstrates that the reduction of VPS50 impairs synaptic transmission in cultured neurons where these deficiencies can be recovered by artificially acidifying synaptic vesicles. *In vitro* electrophysiological recordings using hippocampal slices, show severe deficit in LTP formation. The strong decrease in the frequency of synaptic events observed in VPS50 mKO synapses suggests an increase in the exocytosis of empty or partially filled vesicle. Consistently, significant effects were observed with high-frequency stimulation, a condition in which one might detect the release of partially filled vesicles. These observations strongly support, that the total content rather than vesicle refilling could be account for the synaptic deficit observed at hippocampal VPS50 mKO synapse. Ultimately, these synaptic transmission defects impair memory formation in animals, linking the deficits in synaptic vesicle acidification and/or synaptic filling to complex cognitive behaviors such as learning and memory formation.

In summary, our data strongly support the role of VPS50 in regulating synaptic transmission by facilitating the recruitment of the V-ATPase pump to synaptic vesicles, thereby enabling vesicle acidification and modulating vesicle content. These functional alterations have significant implications, as VPS50 mKO mice exhibit cognitive impairment. Future studies will investigate additional complex behaviors to provide a comprehensive understanding of the behavioral consequences associated with VPS50 mKO including its associated to ASD.

## Acknowledgments

This work was supported by the Chilean government through ANID FONDECYT Iniciacion #11180540 (FJB), PAI #77180077 (FJB), FONDECYT Regular # 1220480 (G.A), #1201848 (AEC), UNAB DI-02-22/REG (FJB), Fondequip # EQM160154 (A.E.C) and by ANID Millennium Science Initiative Program (P09-022F to A.E.C). A grant from the Simons Foundation to the MIT Simons Center for the Social Brain (to H. Robert Horvitz and Martha Constantine-Paton), NIH grant R01GM024663 (to H. Robert Horvitz), NIH grant R01-EY014420 (to M.C.P.). H.R.H. is the David H. Koch Professor of Biology at MIT and an Investigator of the Howard Hughes Medical Institute. CA-G was supported by PhD fellowship from ANID #21201603.

## Figure legends

**Supplementary Figure 1.**
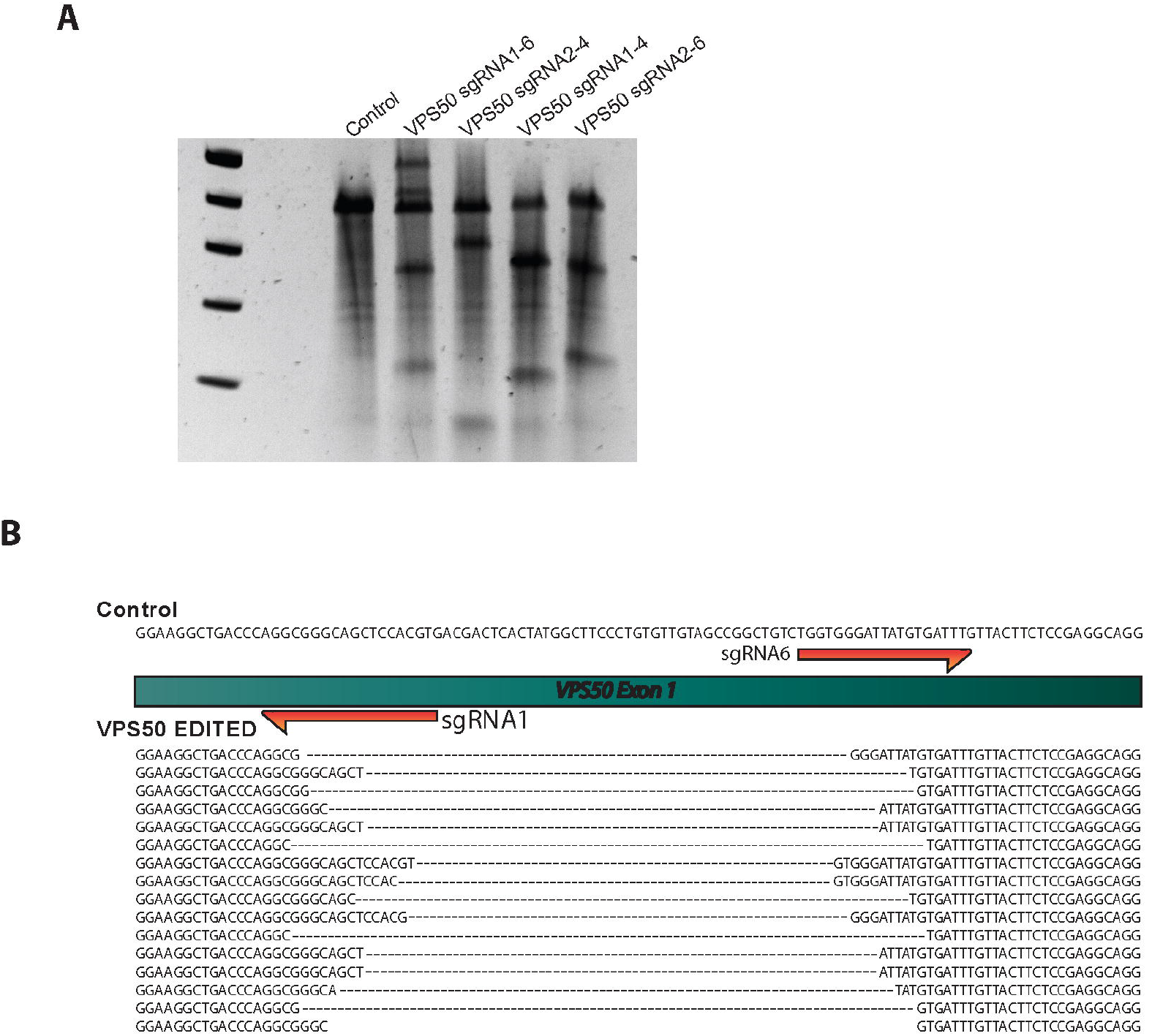
Cortical neurons were infected at 3 DIV with AVV coding for TdTomato as control and a combination of sgRNAs (1-6 / 2-4 / 1-4 / 2-6) targeting Vps50, as shown in figure. 10 days later genomic DNA was extracted, and Surveyor assay performed (A). Surveyor assay for sgRNA combinations (B) Vps50 locus specific sequencing of edited genomic DNA using VPS50 mKO.

**Supplementary Figure 2.**
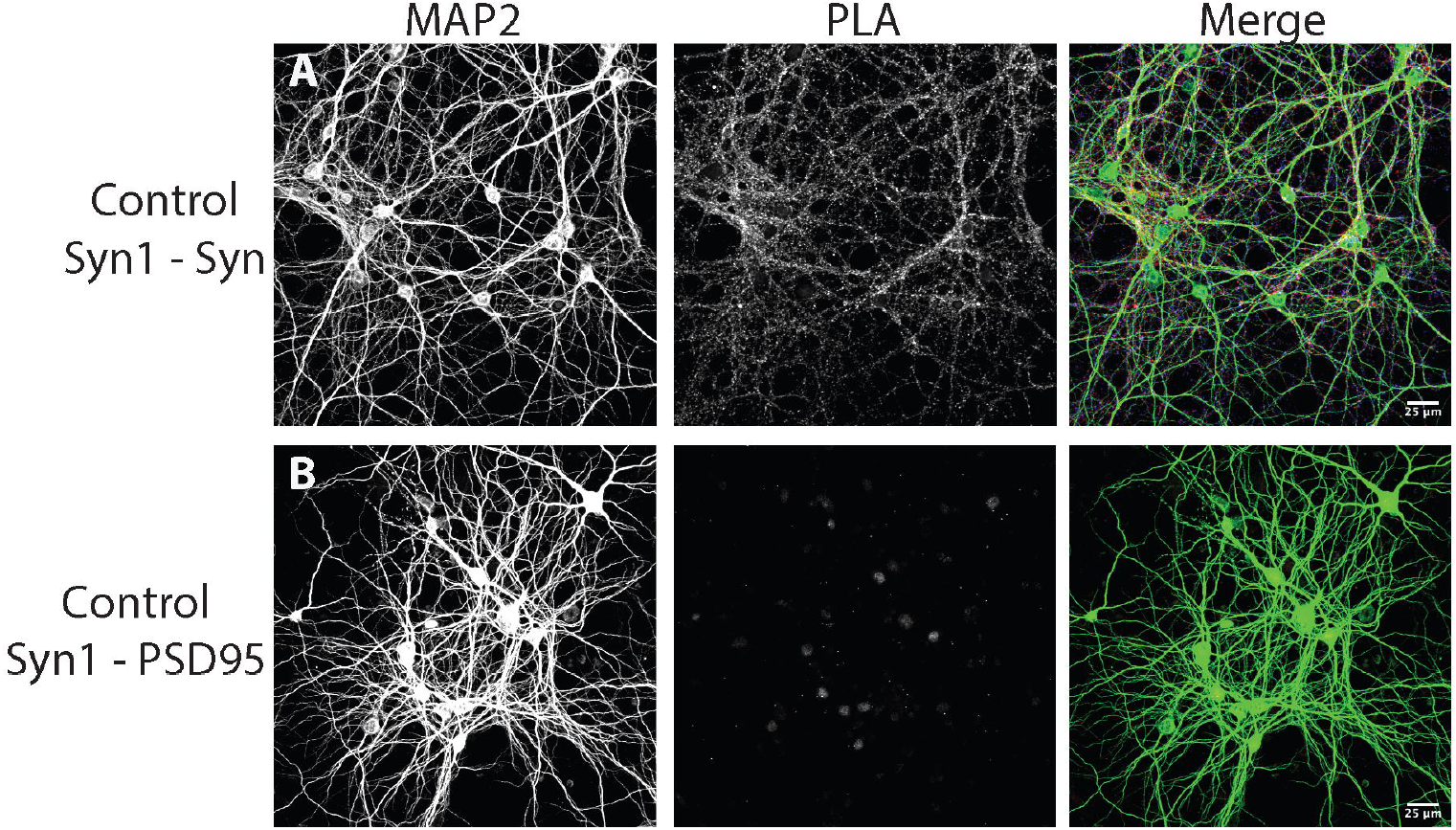
Proximity ligation assay (PLA) assay controls. PLA was performed using specific antibodies for (A) Synapsin1-Synaptosophysin (pre-pre synaptic) or (B) Synapsin1-PSD95 (pre-post synaptic), as shown in figure. PLA signal is only observed in Synapsin1-Synaptosophysin pair where the two proteins are in close proximity to each other. Scale bar, 25 um.

**Supplementary Figure 3.**
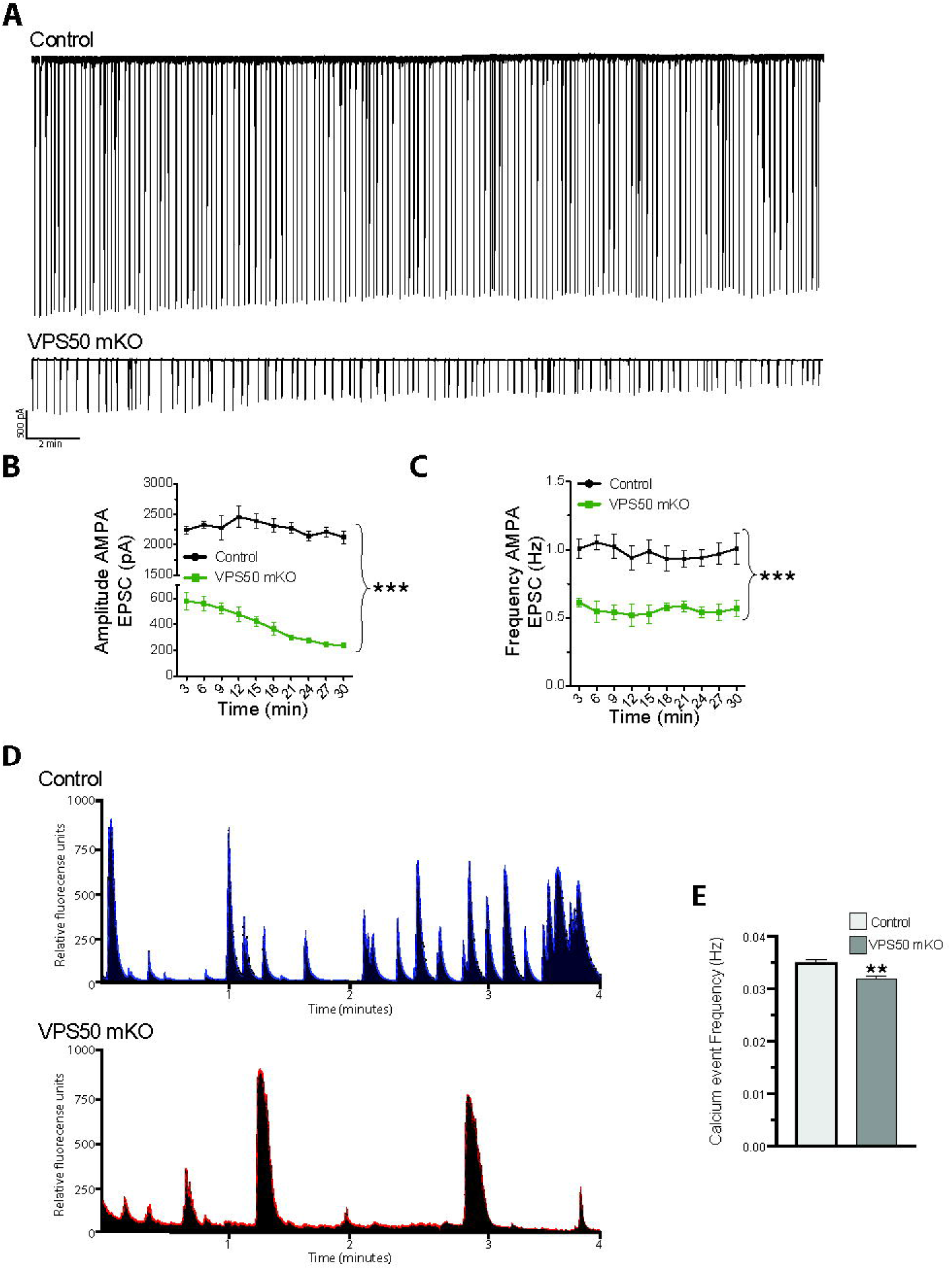
VPS50 mKO neurons show deficits in synaptic transmission. (A) Representative traces of AMPA mediated EPSCs for 30 minutes of Control and VPS50 mKO neurons. VPS50 mKO neurons show a robust reduction in both Amplitude (B) and Frequency (C) of AMPA EPSCs. (D) Cortical neurons were co-transduced with GCaMP7 to measure calcium events by changes in fluorescence over time. A significant reduction is observed in VPS50 mKO neurons compared to control. Control n=1338; VPS50 mKO n=1256. **p<0.01, ***p<0.001.

